# Zika virus persistence in the male macaque reproductive tract

**DOI:** 10.1101/2022.03.03.482872

**Authors:** Erin E. Ball, Patricia Pesavento, Koen K. A. Van Rompay, M. Kevin Keel, Anil Singapuri, Jose P. Gomez-Vazquez, Dawn M. Dudley, David H. O’Connnor, Meghan E. Breitbach, Nicholas J. Maness, Blake Schouest, Antonito Panganiban, Lark L. Coffey

## Abstract

Zika virus (ZIKV) is unique among mosquito-borne flaviviruses in that it is also vertically and sexually transmitted by humans. The male reproductive tract is thought to be a ZIKV reservoir; however, the reported magnitude and duration of viral persistence in male genital tissues varies widely in humans and non-human primate models. ZIKV tissue and cellular tropism and potential effects on male fertility also remain unclear. The objective of this study was to resolve these questions by analyzing archived genital tissues from 51 ZIKV-inoculated male macaques and correlating data on plasma viral kinetics, tissue tropism, and ZIKV-induced pathological changes in the reproductive tract. We hypothesized that ZIKV would persist in the male macaque genital tract for longer than there was detectable viremia, where it would localize to germ and epithelial cells and associate with lesions. We detected ZIKV RNA and infectious virus in testis, epididymis, seminal vesicle, and prostate gland. In contrast to prepubertal males, sexually mature macaques were significantly more likely to harbor persistent ZIKV RNA or infectious virus somewhere in the genital tract, with detection as late as 60 days post-inoculation. ZIKV RNA localized primarily to testicular stem cells/sperm precursors and epithelial cells, including Sertoli cells, epididymal duct epithelium, and glandular epithelia of the seminal vesicle and prostate gland. ZIKV infection was associated with microscopic evidence of inflammation in the epididymis and prostate gland of sexually mature males, which could have significant effects on male fertility. The findings from this study increase our understanding of persistent ZIKV infection which can inform risk of sexual transmission during assisted reproductive therapies as well as potential impacts on male fertility.

**Author Summary:** Zika virus (ZIKV) spread since 2015 led to establishment of urban epidemic cycles involving humans and *Aedes* mosquitoes. ZIKV is also sexually and vertically transmitted and causes congenital Zika syndrome. Together, these features show that ZIKV poses significant global public health risks. By virtue of similar reproductive anatomy and physiology to humans, macaques serve as a useful model for ZIKV infection. However, macaque studies to date have been limited by small sample size, typically 1 to 5 animals. Although mounting evidence identifies the male reproductive tract as a significant ZIKV reservoir, data regarding the duration of ZIKV persistence, potential for sexual transmission, and male genitourinary sequelae remain sparse. Here, we analyzed archived genital tissues from more than 50 ZIKV-inoculated male macaques. Our results show that ZIKV can persist in the male macaque reproductive tract after the resolution of viremia, with virus localization to sperm precursors and epithelial cells, and microscopic evidence of inflammation in the epididymis and prostate gland. Additionally, we show that freezing is not a viable method of destroying infectious ZIKV. Our findings help explain cases of sexual transmission of ZIKV in humans, which also carries a risk for transmission via assisted fertility procedures, even after resolution of detectable viremia.

## Introduction

Mosquito-borne Zika virus (ZIKV) rapidly emerged into urban areas in 2007 (1), initiating epidemics in the South Pacific and the Americas since 2015 and resulting in over 40,000 ZIKV cases in the U.S. and its territories (2). Coupled with this swift global vector-borne spread is the capacity for ZIKV to also be sexually and vertically transmitted, although there are currently no diagnostic mechanisms to distinguish sexually transmitted ZIKV from mosquito-acquired infection (3). Cases of sexual ZIKV transmission are likely underreported owing to a high number of asymptomatic individuals, where passive surveillance shows that 4 out of 5 infections do not produce disease (4). Recent evidence indicates that sexual transmission of ZIKV may be responsible for a significant number of infections and could also serve as mechanism for introducing ZIKV to non-endemic regions lacking mosquito-human-mosquito transmission (5–7).

There is strong evidence that the male reproductive tract serves as an important ZIKV reservoir. Infectious ZIKV and ZIKV RNA have been identified in the semen of symptomatic and asymptomatic men, as well as in vasectomized men, suggesting that, in addition to the testes and epididymis, the virus likely persists in the bulbourethral glands, prostate gland, and/or seminal vesicles (8, 9). However, reported durations of viral persistence in semen and male genital tissues vary widely. Viral RNA has been detected in human semen for up to 370 days after the onset of symptoms, while infectious virus is more short-lived, with positive cultures from semen samples reported for up to 69 days (5, 10). It remains unclear whether there is an association between the magnitude and duration of viremia and genital invasion by ZIKV, viral shedding in semen, and subsequent risk of male-to-female or male-to-male sexual transmission. Genitourinary sequelae in ZIKV infection are not well described, with the exception of hematospermia, prostatitis, and low sperm counts, which are occasionally reported in ZIKV-infected men (8, 9).

While laboratory mice serve as a useful, tractable models of human infectious disease, including ZIKV infection, rhesus and cynomolgus macaques have a closer genetic relationship and reproductive anatomy and physiology comparable to humans, with the same primary and secondary sex organs, similar stages of spermatogenesis, and comparable levels of male sex hormones (11, 12). Thus, extrapolating data from macaque models of ZIKV is a useful mechanism for understanding the pathogenesis of and risk factors associated with human ZIKV infection. *In vivo* viral kinetics, including the length and magnitude of viremia, ZIKV RNA and infectious virus levels within tissues, and tissue tropism, are similar in adult macaques and humans (13–15), validating macaques as a useful model for human ZIKV infection. Unfortunately, long-term data regarding the duration and magnitude of ZIKV in the male genital tract, as well as tropism for specific cell types, are sparse in non-human primates (NHP). ZIKV RNA in rhesus and cynomolgus macaques has been detected in the semen for up to 28 days post-inoculation (DPI), after the resolution of viremia (16). ZIKV RNA has also been detected in the testes (16, 17), prostate gland and seminal vesicles (16, 18) of 6 macaques from 4 to 35 DPI. These time points represent the end of studies, so viral persistence in male macaque genital tissues and shedding in semen may be more prolonged than suggested by published data.

Although significant lesions and evidence of infertility are infrequently reported in NHP, mouse models have variously demonstrated ZIKV-induced orchitis with seminiferous tubule necrosis, testicular atrophy, oligospermia, viral tropism for spermatogenic precursors and Sertoli cells (19–21). Overall, the duration of ZIKV persistence in the male reproductive tract, specific viral tissue and cellular tropisms, and potential effects of ZIKV on male genitourinary symptoms remain unclear in both humans and animal models. Further information on ZIKV persistence in the male reproductive tract can improve guidelines regarding risks of male sexual transmission and infertility. These gaps in knowledge also affect the field of assisted reproductive technology (ART). There are no documented instances of ZIKV transmission due to assisted fertility procedures; however, ZIKV transmission via sperm, oocytes, or embryos is theoretically possible (22). Notably, there is one documented case of congenital Zika syndrome in a fetus associated with sexual transmission from an asymptomatic man with a history of travel to a ZIKV endemic area to his pregnant wife (23).

Based on published data (14,16,18–21), we hypothesized that ZIKV would persist in the male macaque genital tract for longer than detectable viremia, where it would localize to germ cells and epithelial cells and associate with lesions. We used archived reproductive tissues from 51 ZIKV-inoculated male macaques from past collaborative research projects at 4 National Primate Research Centers (NPRC) for this study. These animals, aged 2 to 15 years old, were each inoculated once or multiple times with different doses, using varied routes and strains of ZIKV. They were euthanized and necropsied at times ranging from 1 to 60 DPI. Using tissues from these animals, we quantified ZIKV RNA and infectious virus in genital tissues using qRT-PCR and plaque assays, localized ZIKV RNA to specific cell types using *in-situ* hybridization (ISH), and evaluated histomorphology in testes, epididymis, seminal vesicle, and prostate gland. Our results suggest that the male macaque reproductive tract indeed serves as a reservoir for ZIKV, where the epididymis and seminal vesicle are most likely to harbor virus. We further demonstrate that ZIKV RNA localizes primarily to stem cells (spermatogonia), sperm precursors (1° and 2° spermatocytes) and various epithelial cells, including Sertoli cells, epididymal duct epithelium, and glandular epithelium of the seminal vesicle and prostate gland. Finally, we show that ZIKV infection is associated with microscopic lesions in the epididymis and prostate gland, which could have significant effects on male fertility.

## Results

### Samples from 51 experimentally ZIKV inoculated male macaques were used to evaluate ZIKV tropism and disease in the male reproductive tract

Archived reproductive tissues and fluids from previous research projects, including testes, epididymis, seminal vesicle, prostate gland and/or semen from 51 ZIKV-inoculated and 8 uninfected male rhesus (*Macaca mulatta*) and cynomolgus (*Macaca fascicularis*) macaques ranging from 2 to 15 years old were provided by the California (CNPRC), Tulane (TNPRC), Wisconsin (WNPRC), and Washington (WaNPRC) National Primate Research Centers (**Table 1**). All uninfected control tissues were from rhesus macaques (N = 7 from CNPRC, N = 1 from WaNPRC). Of the ZIKV-inoculated animals, 5 were cynomolgus macaques from TNPRC, while the remaining 46 were rhesus macaques from CNPRC, WNPRC and TNPRC. Twenty-two animals (all rhesus macaques) were inoculated intravenously (IV) and 28 (including the 5 cynomolgus macaques) were inoculated subcutaneously (SC) with Brazilian (N = 20), Puerto Rican (N = 23), or French Polynesian (N = 2 [semen samples]) strains of ZIKV and necropsied between 1 and 60 DPI. Five ZIKV-inoculated rhesus macaques were inoculated with plasma from a ZIKV-infected human for which specific strain information was not available, and the duration of infection was not available for 2 rhesus macaques for which frozen semen was the only submitted sample. Animals lacking specific metadata were not included in statistical analyses. Among macaques from CNPRC and TNPRC, 4 were immune suppressed via CD8+-cell depletion to assess the impact of these cells on acute ZIKV infection immediately prior to (N = 2 cynomolgus macaques) or 4 weeks after (N = 2 rhesus macaques) ZIKV inoculation. Seven sexually immature rhesus macaques received anti-ZIKV antibody prior to inoculation (5 of 7 animals for which viremia data was available had reduced or delayed viremia), while 9 animals that failed to become viremic upon initial inoculation were reinoculated with a higher dose to ensure infection. Plasma and serum and or viremia data was available from only a subset of the animals, N = 36. There was a single vasectomized from CNPRC and a single splenectomized cynomolgus macaque from TNPRC.

**Table 1:**
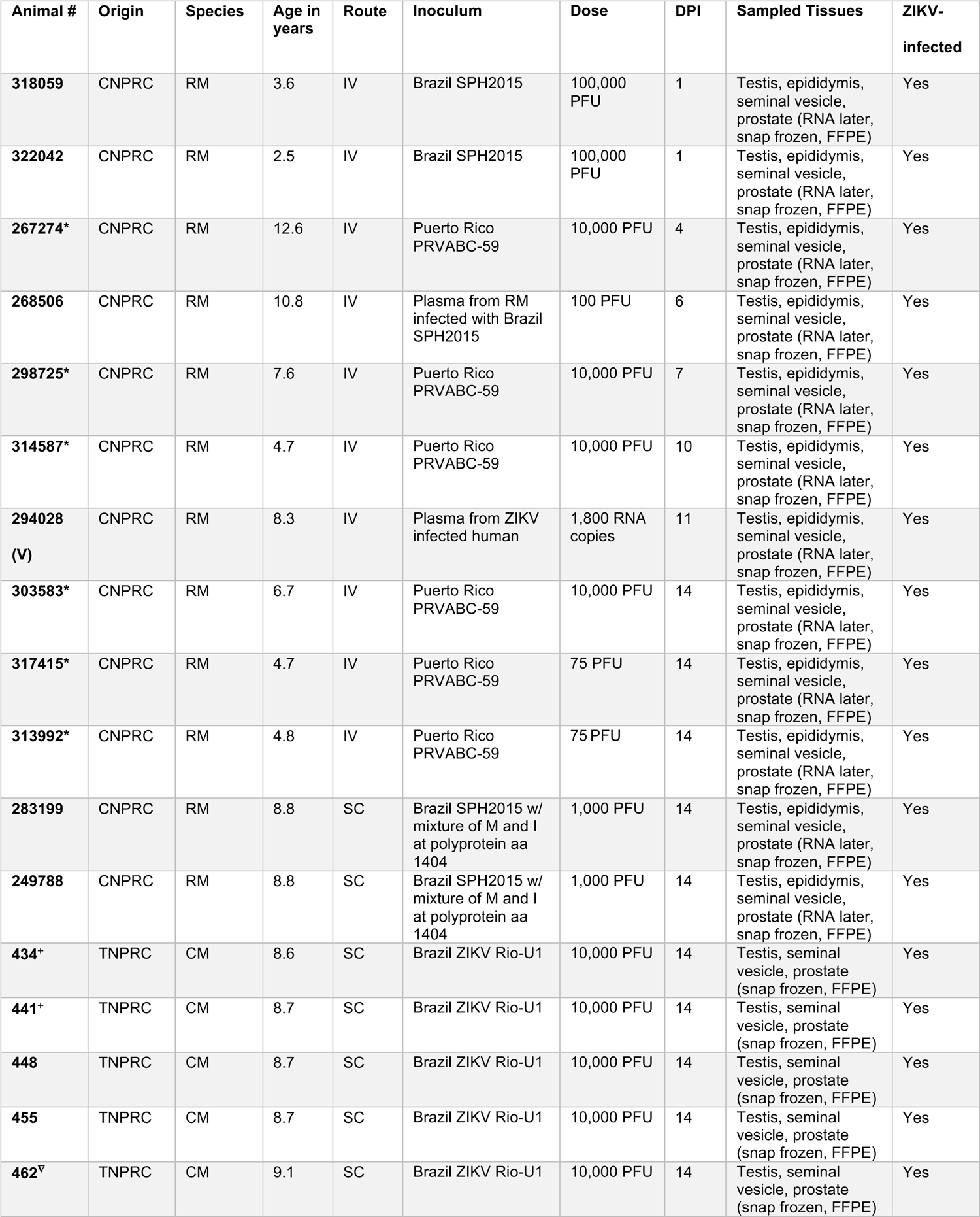
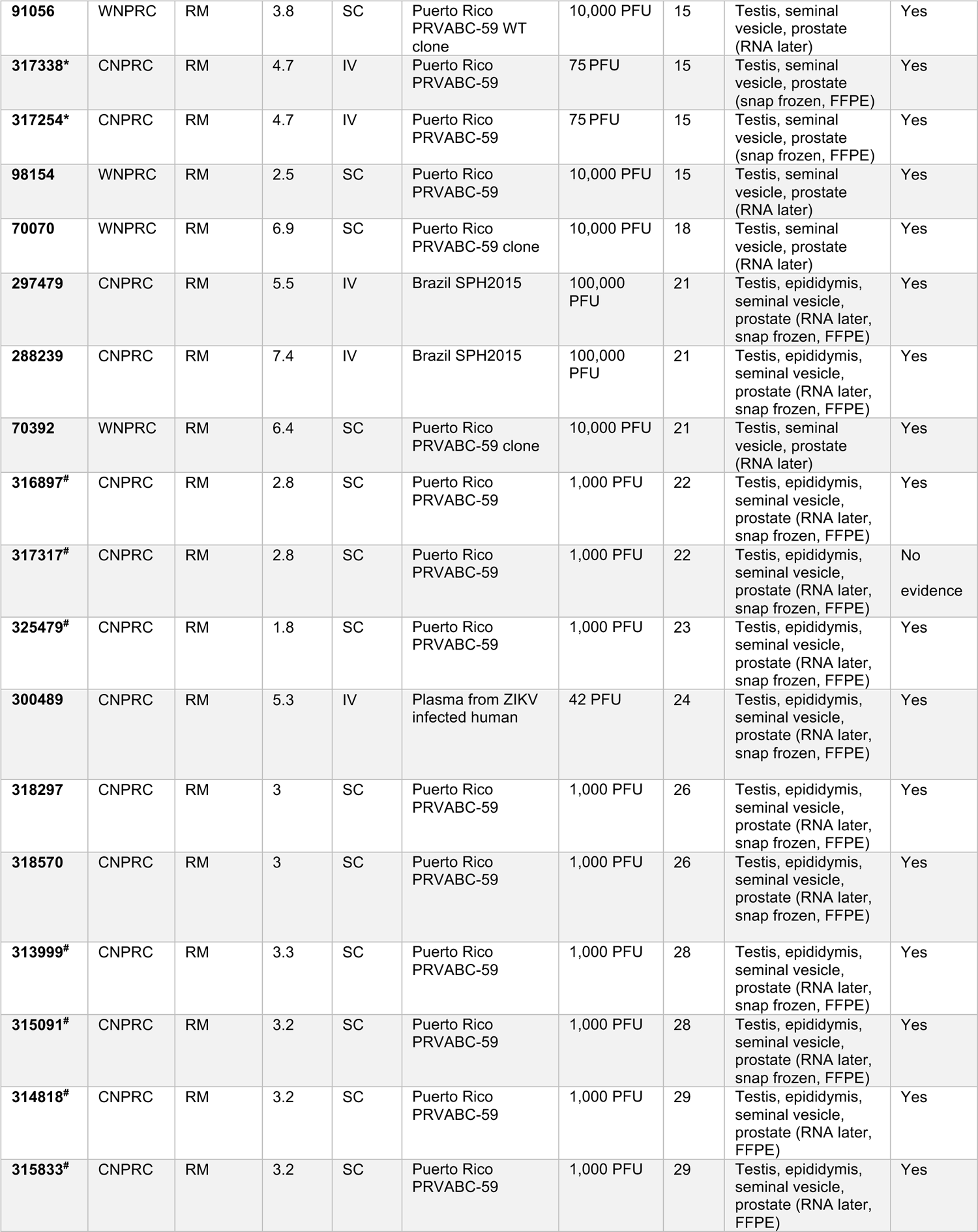
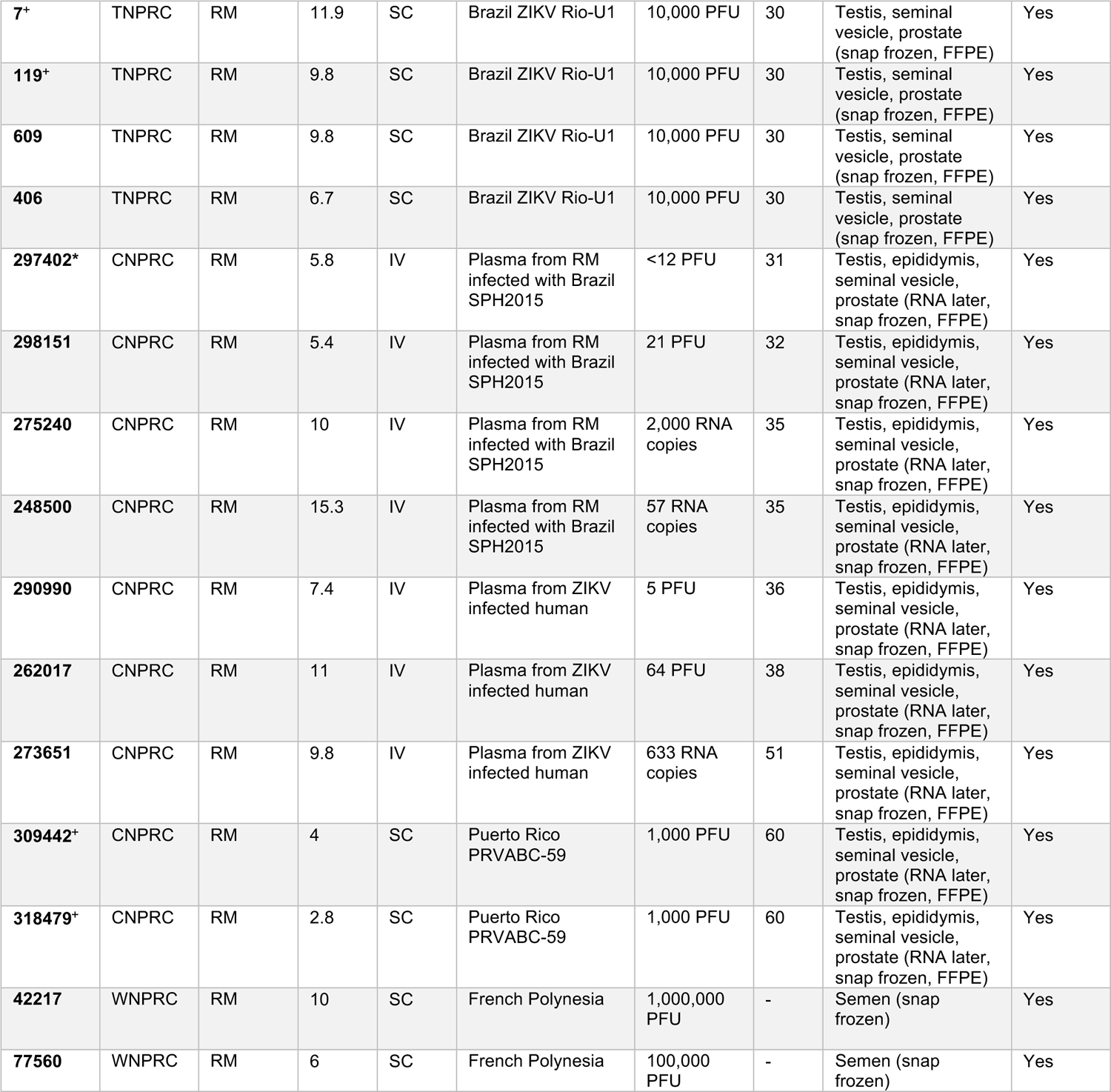
Male macaques and the ZIKV treatments used on animals in this study. A macaque was considered ZIKV-infected if ZIKV RNA was detected in any fluid or tissue. * Reinoculated animals; (V) vasectomized animal; ^#^ received anti-ZIKV antibody prior to inoculation; ^+^ immune suppressed animals; ^∇^splenectomized animal; (semen) frozen semen was the only submitted sample; ‘-’data not available; RM= rhesus macaque; CM is cynomolgus macaque; IV is intravenous; SC is subcutaneous; ZIKV is Zika virus; PFU is plaque forming units; DPI is days post inoculation the animal was euthanized. For ZIKV strains, Brazil SPH2015 is GenBank accession KU321639.1, Brazil ZIKV Rio-U1 is GenBank accession KU926309, Puerto Rico PRVABC-59 is GenBank accession KU501215.1, PRVABC-59 clone is PRVABC-59 clone virus (pjW236-C1 P0), PRVABC-59 WT clone is PRVABC-59 clone virus (pW232-WT P0), French Polynesia Zika virus/*H.sapiens*-tc/FRA/2013/FrenchPolynesia/3328 is GenBank accession KJ776791.2. CNPRC is California National Primate Research Center; TNPRC is Tulane National Primate Research Center; WNPRC is Wisconsin National Primate Research Center; FFPE is formalin fixed paraffin embedded tissues; w/ is with, aa is amino acid.

### A subset of differing experimental conditions exert significant effects on ZIKV detection and histopathologic lesion severity in male macaque reproductive tissues

We first assessed the effects of different experimental conditions, including the frequency and route of ZIKV inoculation, viral strain, dose, macaque species, administration of anti-ZIKV antibody, and immune suppression, on whether ZIKV RNA, infectious ZIKV, and microscopic lesions were detected in male genital tissues. Viremia and detection of ZIKV in male reproductive tissue were significantly associated with the dose and route (IV versus SC) of inoculation (Supplemental Table 1). A higher ZIKV inoculation dose resulted in higher peak viremia (Kruskal Wallis, p = 0.032). IV inoculation was also associated with significantly higher peak viremia (p = 0.02; 0.21, 1.89 95% confidence interval [CI]) than SC inoculation, and was 8.71 (1.52, 94.17 95% CI) times more likely than SC inoculation to result in ZIKV detection somewhere in the male reproductive tract; 13.5 (2.23, 115.66 95% CI) times more likely in the epididymis, and 6.11 (1.41, 31.35 95% CI) more likely in the seminal vesicle. Route of inoculation was not significantly associated with the likelihood of detecting ZIKV in the prostate or testis. Likewise, the IV route of inoculation was associated with a higher histology severity score in the epididymis (0.36, 1.70 95% CI) and prostate gland (0.01, 1.10 95% CI). Of the 7 animals treated with anti-ZIKV antibody, all 5 animals for whom plasma was available had detectable viremia, although peak viremia was delayed and reduced in magnitude when compared to untreated controls. Administration of anti-ZIKV antibody was also associated with a decreased likelihood of detecting of ZIKV RNA within the epididymis (Fisher’s exact, p = 0.01) and significantly lower histology severity scores within the prostate gland (simple logistic regression, p < 0.0001). Reinoculation with a second dose of ZIKV after no infection was detected as a result of the first inoculation resulted in a higher viremia area under the curve (AUC; p = 0.002; 10.68, 43.94 95% CI) and significantly affected histology severity scores in sexually mature macaques, increasing the epididymis score by 1.62 (0.88, 2.6 95% CI) and the prostate gland score by 0.95 (0.30, 1.61 95% CI). Immune-suppression by CD8 T-cell depletion and viral inoculum strain (Brazil vs. Puerto Rico) did not significantly affect presence of ZIKV RNA or histology severity scores within male reproductive tissues. The limited sample size for cynomolgus (5 animals) versus rhesus (46 animals) macaques precluded statistical assessments of associations between species and the dependent variables. The experimental conditions exerting a significant effect on the dependent variables (including the presence of ZIKV RNA, infectious ZIKV and microscopic lesion severity in genital tissues) were subsequently incorporated into the appropriate ordered logistic regression or linear model.

### Peak viremia magnitude and viremia area under the curve (AUC) are significantly higher in sexually mature versus immature ZIKV-inoculated male macaques

A macaque was considered ZIKV-infected if viremia or ZIKV RNA was detected in any other fluid or tissue besides serum or plasma (**Table 1**). Viremia was assessed from ZIKV RNA in sera or plasma in 36 of the adult male macaques and was detected in 34 of the 36 animals. Viremia in macaques, excluding animals that were pre-treated with anti-ZIKV antibody that showed delayed and reduced magnitudes, typically lasted for 4 to 14 DPI with a peak between 4 and 8 log_10_ RNA copies/mL and occurring, on average, at 4 to 5 DPI (**Figure 1**). Of the 15 animals for which viremia data was unavailable and the one animal without detectable viremia, ZIKV RNA was detected in at least one genital tissue or fluid (including semen) for 14 animals, confirming ZIKV infection. The single remaining rhesus macaque did not have detectable viral RNA in any fluid or tissue, so ZIKV infection could not be definitively confirmed. Peak viremia magnitude (ordered logistic regression model, p = 0.002) and AUC (ordered logistic regression model, p = 0.03) were significantly higher in sexually mature ZIKV-inoculated male macaques when compared to pre-pubertal males. Higher peak viremia correlated with ZIKV RNA detection in the epididymis (ordered logistic regression model, p = 0.042), but otherwise, viremia kinetics did not correlate with ZIKV RNA, infectious ZIKV levels (Kruskal Wallis, all remaining p values > 0.05), or histologic lesion severity in male genital tissues (Spearman’s rank correlation, all p values > 0.05) (**Supplemental Table 1**).

**Figure 1.**
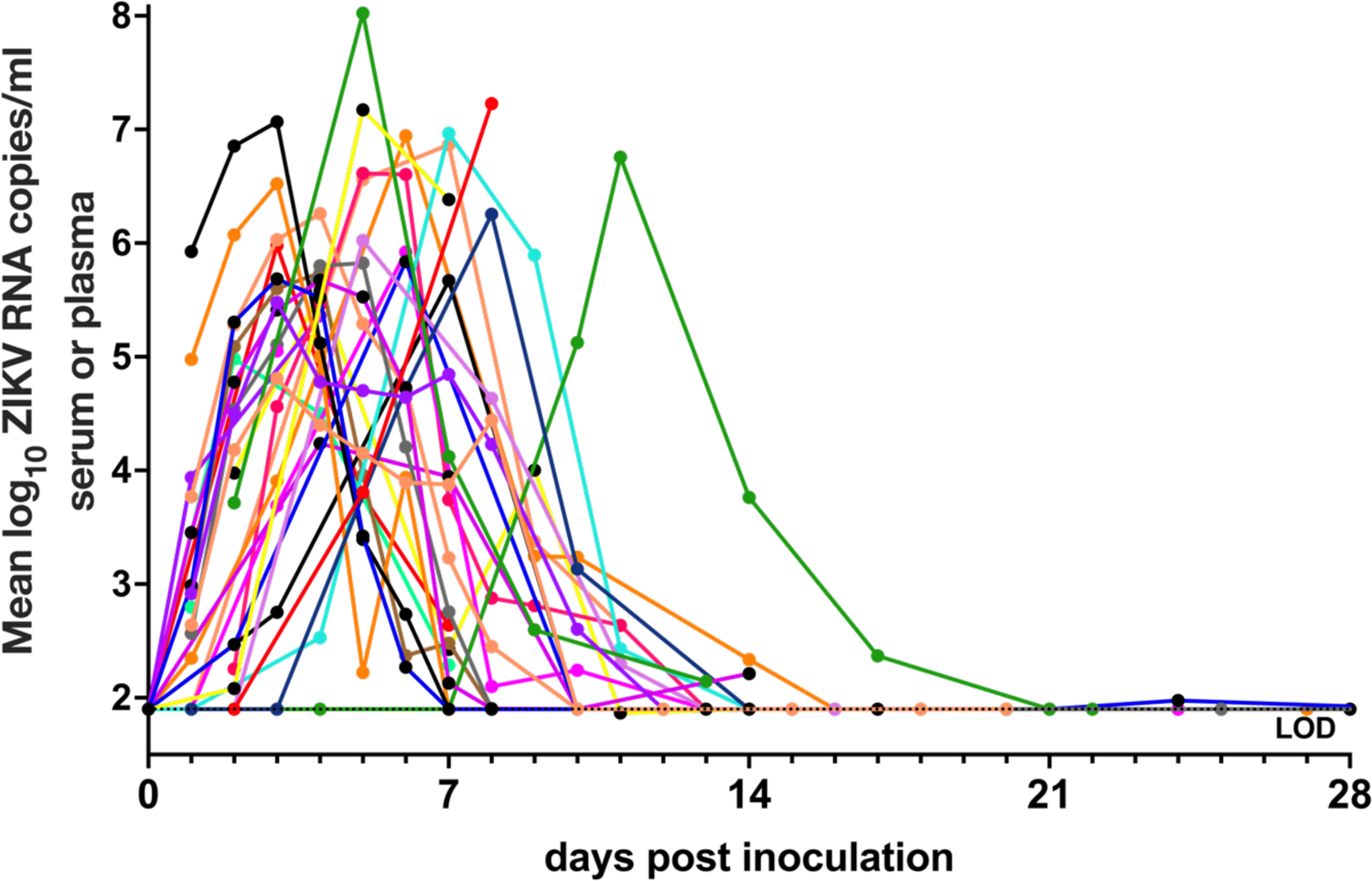
Zika virus viremia in adult male macaques lasts for 4 to 25 days post-inoculation with a peak from 4 - 8 log_10_ RNA copies/mL at 4 - 5 days post-inoculation (DPI). ZIKV RNA levels in serum or plasma, reported as mean log_10_ RNA copies/ml and assayed in triplicate. Each line/symbol represents an individual macaque (N = 36). Viremia data was not available for each of the 51 animals. Rhesus macaques pre-treated with anti-ZIKV antibody were not included in calculations of average viremia duration and magnitude and are not shown on this graph. The dotted line denotes the average limit of detection (LOD), 1.9 log_10_ RNA copies/ml.

### ZIKV RNA is detectable in the testis, epididymis, seminal vesicle, prostate gland and/or semen of male macaques and can persist for at least 60 days

Both ZIKV RNA and infectious virus were detected in the male macaque reproductive tract. ZIKV RNA was detected in at least one reproductive tissue or fluid including the testis, epididymis, seminal vesicle, prostate gland, or semen in 34 out of 48 macaques (71%). Overall ZIKV RNA was most frequently detected in the seminal vesicle (46%, 22/48) and epididymis (55%, 21/38), while the prostate gland (32%, 15/46) and testes (31%, 16/51) were less frequently ZIKV RNA positive (**Figure 2A**). The highest absolute magnitude of ZIKV RNA was detected in the epididymis at 31 DPI. ZIKV RNA was detected as early as 1 DPI and late as 60 DPI in the testis. The single vasectomized rhesus macaque had detectable ZIKV RNA in all four genital tissues. Previously, 35 DPI was the longest documented duration of ZIKV RNA in male macaque genital tissues (18).

**Figure 2.**
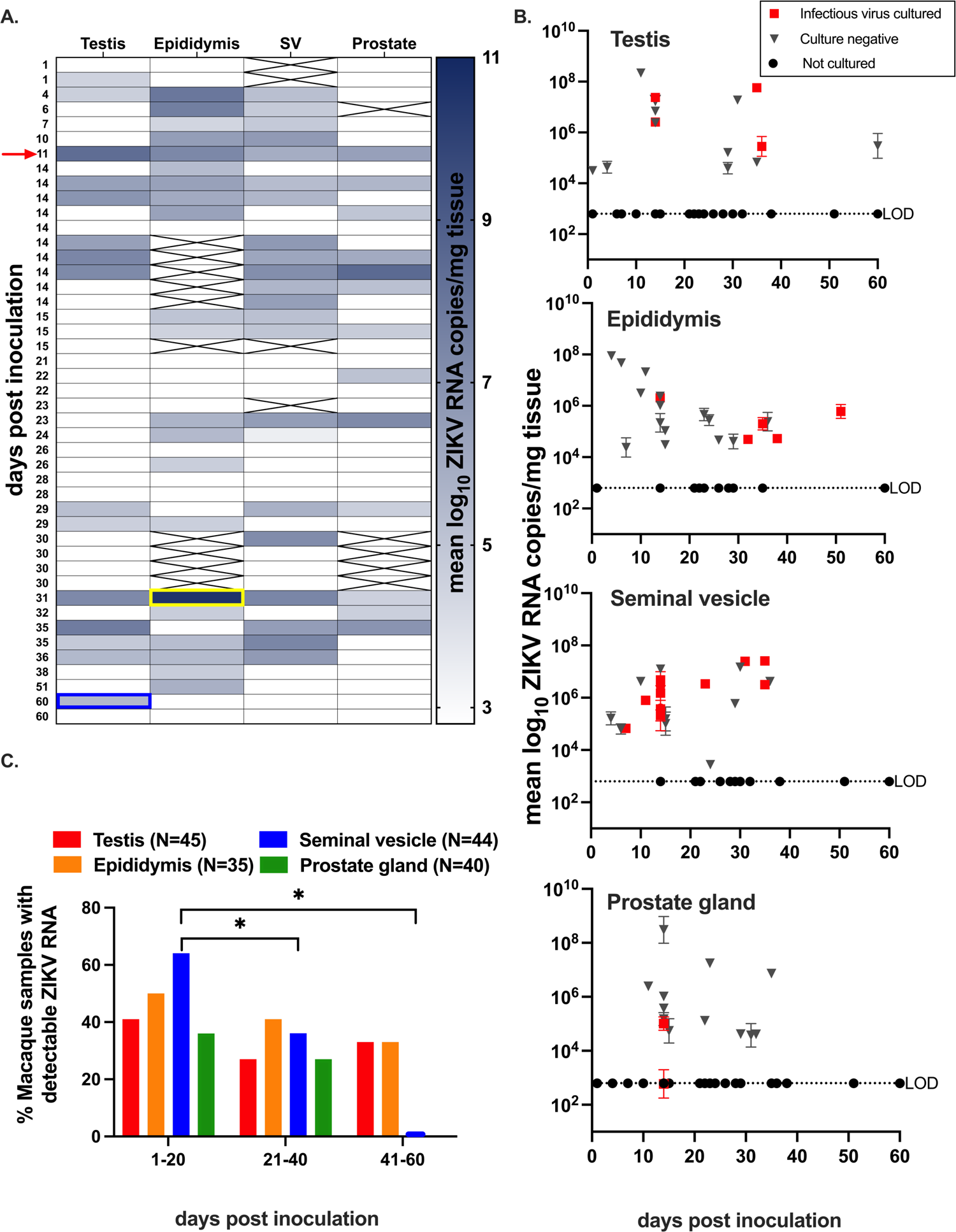
ZIKV RNA and infectious virus are detected in testis, epididymis, seminal vesicle, and prostate gland of male macaques. **A.)** Heatmap categorizing ZIKV RNA by male macaque genital tissue type. The color intensity correlates with the magnitude of ZIKV RNA detected by qRT-PCR and is shown as mean log_10_ RNA copies/mg tissue. A full complement of genital tissues was not collected for every animal and tissues that were not available are crossed out. The highest absolute magnitude was detected in the epididymis (yellow box) and the latest day post inoculation of ZIKV detection occurred in the testis (blue box). There was one vasectomized macaque who had detectable ZIKV RNA in all four genital tissues (red arrow). SV is seminal vesicle. **B.**) ZIKV RNA levels in male macaque genital tissues, reported as mean log_10_ RNA copies/mg tissue and assayed in triplicate with error bars showing standard deviations. Each symbol represents an individual macaque. ZIKV RNA positive samples that also contained detectable infectious ZIKV are denoted with red squares. Black triangles indicate PCR-positive samples which were negative for infectious virus. Black circles denote PCR-negative samples that were not cultured. Titers ranged from 1 - 10 PFU/mg tissue. Average LODs for ZIKV RNA and infectious virus were 2.8 log_10_ RNA copies/mg and 0.4 PFU/mg tissue, respectively. **C.** ZIKV RNA was significantly more likely to be detected in the seminal vesicle (blue bars) at 1-20 DPI (vs. 21-40 DPI and 41-60 DPI). Mann-Whitney: ns, p value > 0.05; *, p value < 0.05; ** p value < 0.01; ***, p value < 0.001; ****p value < 0.0001.

To identify the presence of infectious ZIKV in the male reproductive tract, we performed plaque assays on available frozen samples for tissues with detectable ZIKV RNA. Infectious ZIKV was detected in at least one reproductive tissue in 18 out of 27 macaque tissues (67%), as early as 4 DPI, and as late as 50 DPI in the epididymis (**Figure 2B**). Titers ranged from 1 to 10 PFU/mg tissue (data not shown). Infectious ZIKV was cultured most frequently from the seminal vesicle (63%, 12/19), followed by epididymis (27%, 5/18) and testis (30%, 4/13). Infectious ZIKV was rarely cultured from the prostate gland (16%, 2/12). While both available semen samples contained detectable ZIKV RNA, there was no evidence of infectious virus in either sample.

The duration of detection of ZIKV RNA in this study ranged from 1 to 60 DPI and shorter study endpoints did not correlate with detection of ZIKV RNA in all male genital tissues. However, macaques euthanized at earlier times (between 1 and 20 DPI) were significantly more likely to harbor ZIKV RNA in the seminal vesicle when compared to those euthanized between 21 and 40 DPI (Mann-Whitney, p = 0.02) or 41 to 60 DPI (Mann-Whitney, p = 0.03) (**Figure 2C**; **Supplemental Table 2**). Taken together, these data indicate that ZIKV can persist in multiple male genital tissues and most frequently and at the highest magnitude in the epididymis and seminal vesicle for up to 60 DPI, 1 month after the end of detectable viremia.

### *In-situ* hybridization (ISH) of genital tissues from ZIKV-inoculated male macaques demonstrates ZIKV RNA in the testis, epididymis, seminal vesicle, and prostate gland

To visualize specific cellular tropism of ZIKV, we next performed ISH on male macaque genital tissues where ≥ 5 RNA copies/mg tissue were detected (N = 20 sexually mature; N = 2 sexually immature). ISH staining was consistent with qRT-PCR data. All sections of sexually mature testis, seminal vesicle, and prostate gland that contained detectable ZIKV RNA were also ISH-positive, where positivity was identified as red cytoplasmic/peri-nuclear signal. Both sexually immature animals with detectable ZIKV RNA also had ISH positive tissues. The same was generally true for the epididymis; however, epididymal tissue sections from 2 out of 17 examined samples lacked a visible ISH signal while demonstrating detectable ZIKV RNA. This may be a function of the very small tissue sample sizes, rather than a true disparity between the two RNA detection methods. No ISH signal was observed in tissues from non-inoculated macaques.

In the testis, ZIKV RNA was detected primarily within 1° and 2° spermatocytes (**Figure 3A**), spermatogonia (germ cells), and Sertoli cells (modified epithelial cells), with rare signal in interstitial Leydig cells and peri-tubular cells. For the epididymis, seminal vesicle, and prostate gland, ZIKV RNA was detected most frequently within ductal (epididymis) and glandular (seminal vesicle and prostate gland) epithelial cells (**Figure 3B-D**). In all 4 tissues, spindle cells located within interstitial or capsular connective tissue also occasionally harbored ZIKV RNA. While these cells cannot be definitively identified without special stains, they are most consistent with being migrating macrophages, fibroblasts or possibly mesenchymal stem cells based on cellular morphology and anatomic location. Overall, ZIKV demonstrated a tropism for stem-like cells and epithelial cells of the male macaque reproductive tract.

**Figure 3.**
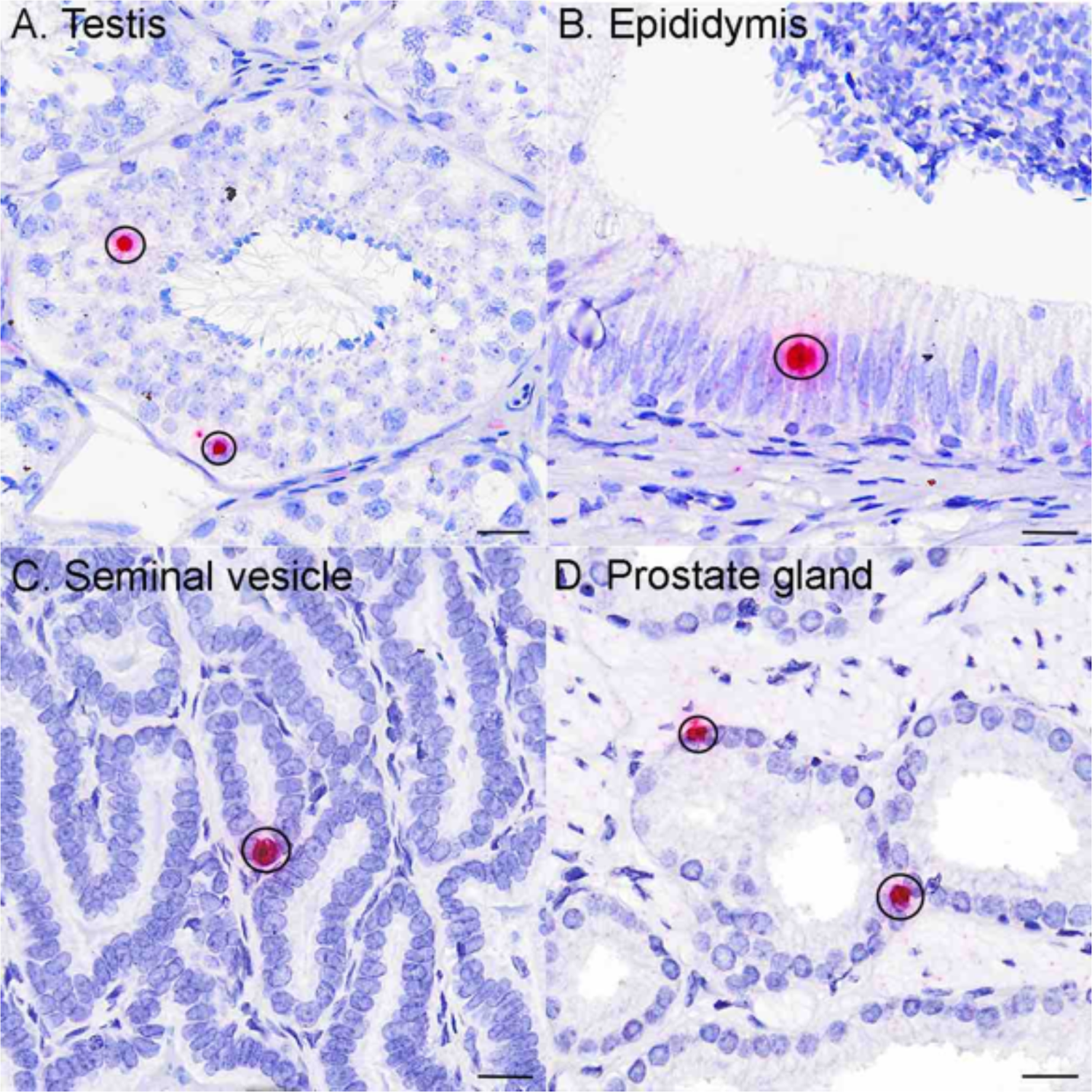
*In-situ* hybridization (ISH) of genital tissues from ZIKV-inoculated male macaques demonstrates ZIKV RNA in the testis, epididymis, seminal vesicle, and prostate gland. Photomicrographs of tissues from ZIKV-inoculated males after ISH where positive cells exhibit an intracytoplasmic/perinuclear red signal (circled). See Figure 7 for normal histology of testis, epididymis, seminal vesicle, and prostate gland. A.) Within the testis ZIKV RNA localized most frequently to sperm precursors, including germ cells, 1°/2° spermatocytes (circled), less frequently to Sertoli cells, and rarely to peritubular spindle cells and Leydig cells (images not shown here). Within the B) epididymis, C) seminal vesicle, and D) prostate gland, ZIKV RNA localized primarily to ductular or glandular epithelial cells (circled). In all 4 tissues, spindle cells (likely fibroblasts or migrating macrophages) located within interstitial or capsular connective tissue, also occasionally harbored ZIKV RNA (images not shown here). No ZIKV staining was observed in tissues from non-inoculated macaques (data not shown). Bar = 20 μm.

### Sexual maturity impacts detection of ZIKV in male macaque reproductive tissues

Male macaques typically reach sexual maturity around 4 years of age (11, 24). Here, 29 out of 36 (81%) sexually mature macaques older than 4 years inoculated with ZIKV had detectable viral RNA in at least one reproductive tissue or fluid, versus just 6 out of 14 (43%) sexually immature macaques. ZIKV RNA was detected more frequently within the reproductive tract of sexually mature male macaques (**Figure 4A**, **Supplemental Table 2**) (ordered logistic regression model, p = 0.004), and specifically in the epididymis (ordered logistic regression model, p < 0.0001) and seminal vesicle (ordered logistic regression model, p = 0.0005) (**Figure 4B**). Similarly, infectious ZIKV was cultured exclusively from male genital tissues of sexually mature macaques. We also assessed effects of sexual maturity on absolute ZIKV RNA magnitude (mean RNA copies/mg tissue); however, due to relatively low Ns and lack of normality, the model resulted in a better fit when overall ZIKV RNA presence or absence in genital tissues was assessed instead. These data demonstrate that sexually mature male macaques are significantly more likely to harbor ZIKV RNA and/or infectious virus somewhere in the reproductive tract.

**Figure 4.**
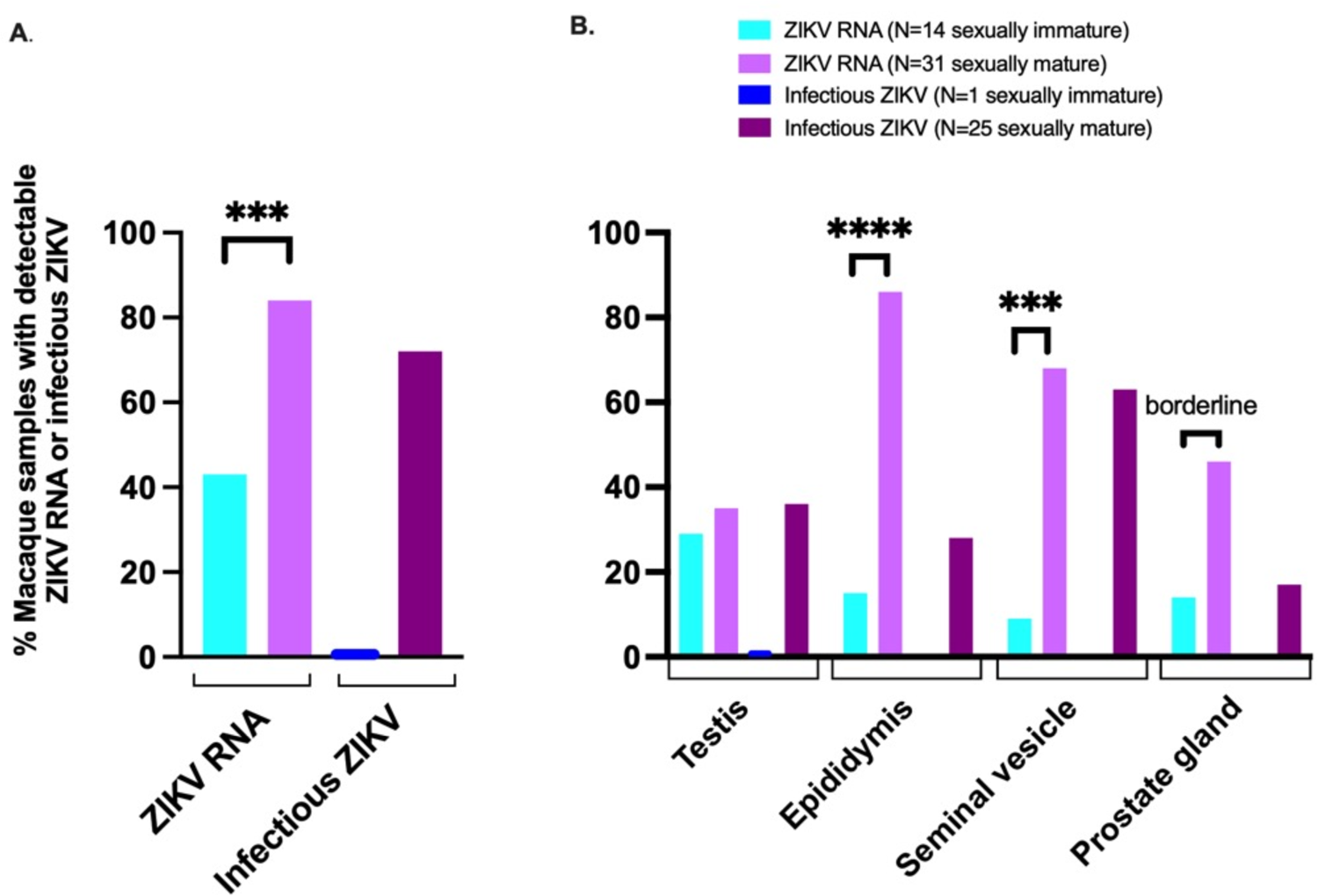
Sexually mature male macaques are more likely to harbor persistent ZIKV RNA and infectious virus in at least one genital tissue. (**A**), in the epididymis and seminal vesicle (**B**). Only one tissue (testis) from a sexually immature macaque had detectable ZIKV RNA and no infectious virus was cultured; statistical tests were not performed on infectious ZIKV data. Sexually immature = 0 - 4 years old; sexually mature = > 4 years old. Ordered logistic regression: ns, p value > 0.08; borderline, p value between 0.05 and 0.08; *, p value ≤ 0.05; ** p value ≤ 0.01; ***, p value ≤ 0.001; ****p value ≤ 0.0001.

### ZIKV inoculation of sexually mature male macaques is associated with microscopic lesions in the epididymis and prostate gland

Overall, histopathologic lesions were relatively uncommon within the reproductive tract of male macaques inoculated with ZIKV. When present, histopathologic lesions in testis, epididymis, seminal vesicle, and prostate gland were scored according to quantitative criteria (**Supplemental Table 3)**. Sexually mature ZIKV-inoculated male macaques had significantly higher severity scores than uninfected, age-matched controls in the epididymis (Mann-Whitney, p = 0.02) and prostate gland (Mann-Whitney, p < 0.001) (**Figure 5A**). Similarly, sexually mature ZIKV-inoculated macaques had significantly more severe microscopic lesions than sexually immature ZIKV-inoculated macaques in the epididymis (linear model, p = 0.02), and prostate gland (linear model, p = 0.0001). No significant lesions were noted in the seminal vesicle. As with ZIKV RNA, macaques euthanized at earlier timepoints (between 1 and 20 DPI) were more likely to have significant microscopic lesions resulting in higher histology scores in the epididymis and prostate gland when compared to those euthanized between 21 and 40 DPI (Mann-Whitney, p = 0.01 and p = 0.02, respectively) or 41 to 60 DPI (Mann-Whitney, p = 0.08 trend and p = 0.05, respectively) (**Figure 5B**, **Supplemental Table 2**).

**Figure 5.**
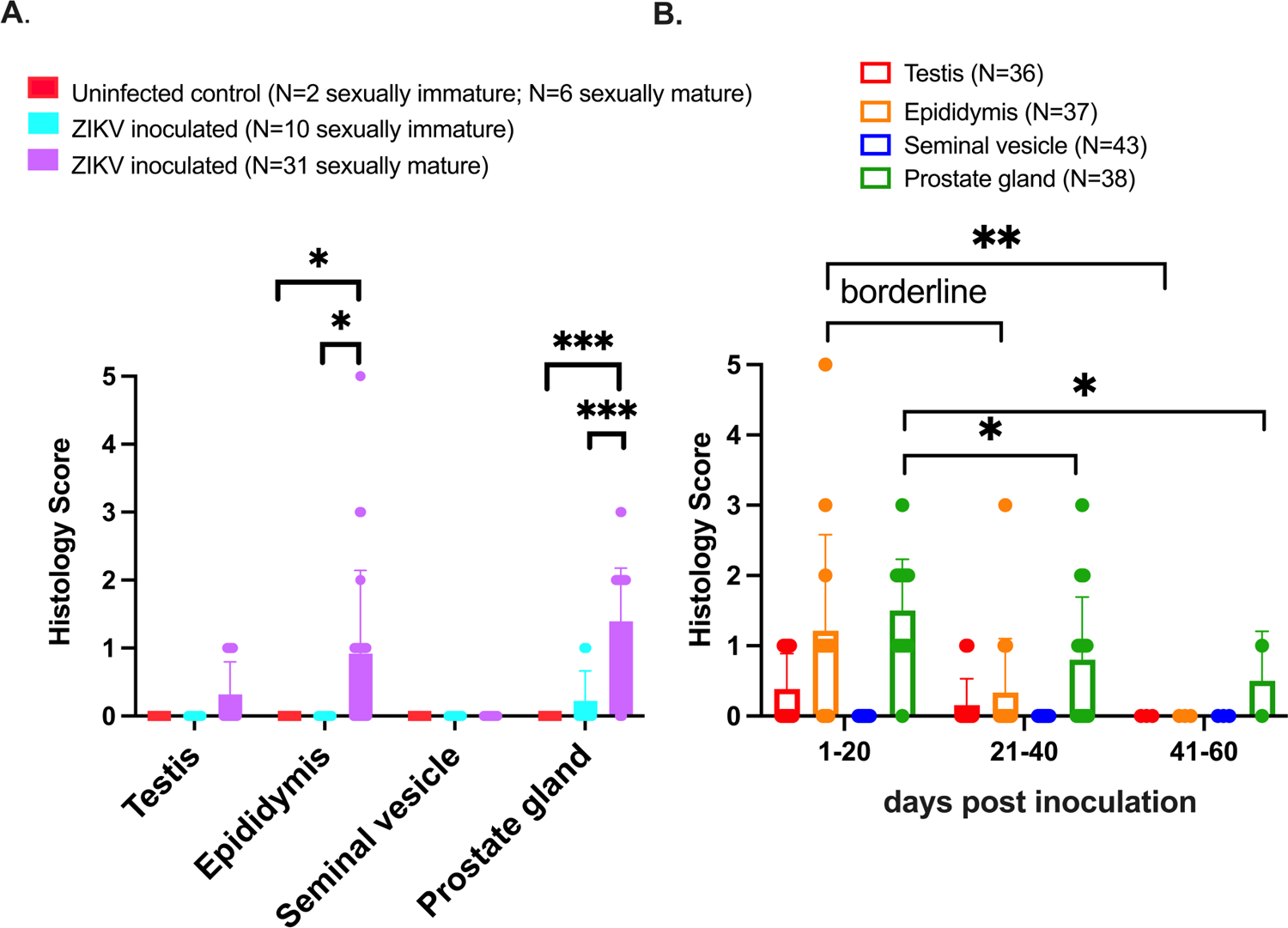
Sexual maturity and duration of ZIKV infection impact histopathologic lesion severity in male macaque reproductive tissues. Macaque genital tissues were scored histologically from 0 (normal) to 5 (markedly abnormal). A.) Pathology severity scores for sexually mature and immature ZIKV-inoculated male macaques and uninfected, age-matched controls B.) Pathologic lesions were more likely to occur between 1 and 20 DPI in the epididymis and prostate gland (vs. 21 - 40 and 41 - 60 DPI). Each dot represents an individual macaque. Bars show the mean histology score, and error bars show standard deviation. Sexually immature = 0 - 4 years old; sexually mature = > 4 years old. Mann-Whitney: ns, p value > 0.08; borderline, p value between 0.05 and 0.08; *, p value ≤ 0.05; ** p value ≤ 0.01; ***, p value ≤ 0.001.

Sporadic epididymal and prostatic inflammation were noted exclusively within sexually mature males. Three out of 25 sexually mature macaques exhibited epididymal microscopic lesions with a histology severity score of ≥ 3. Lesions ranged from mild lymphohistiocytic periductal infiltrates to severe pyogranulomatous epididymitis with duct rupture, multinucleated giant cells containing engulfed spermatozoa, multifocal mineralization, fibroplasia, and sperm stasis with dilated/tortuous epididymal ducts (**Figure 6A, B**). This correlates with the virology data reported above, where macaques with detectable ZIKV RNA in the epididymis exhibited higher histology scores indicative of more severe lesions than those without detectable virus (linear model, p = 0.01, **Supplemental Table 2**).

**Figure 6.**
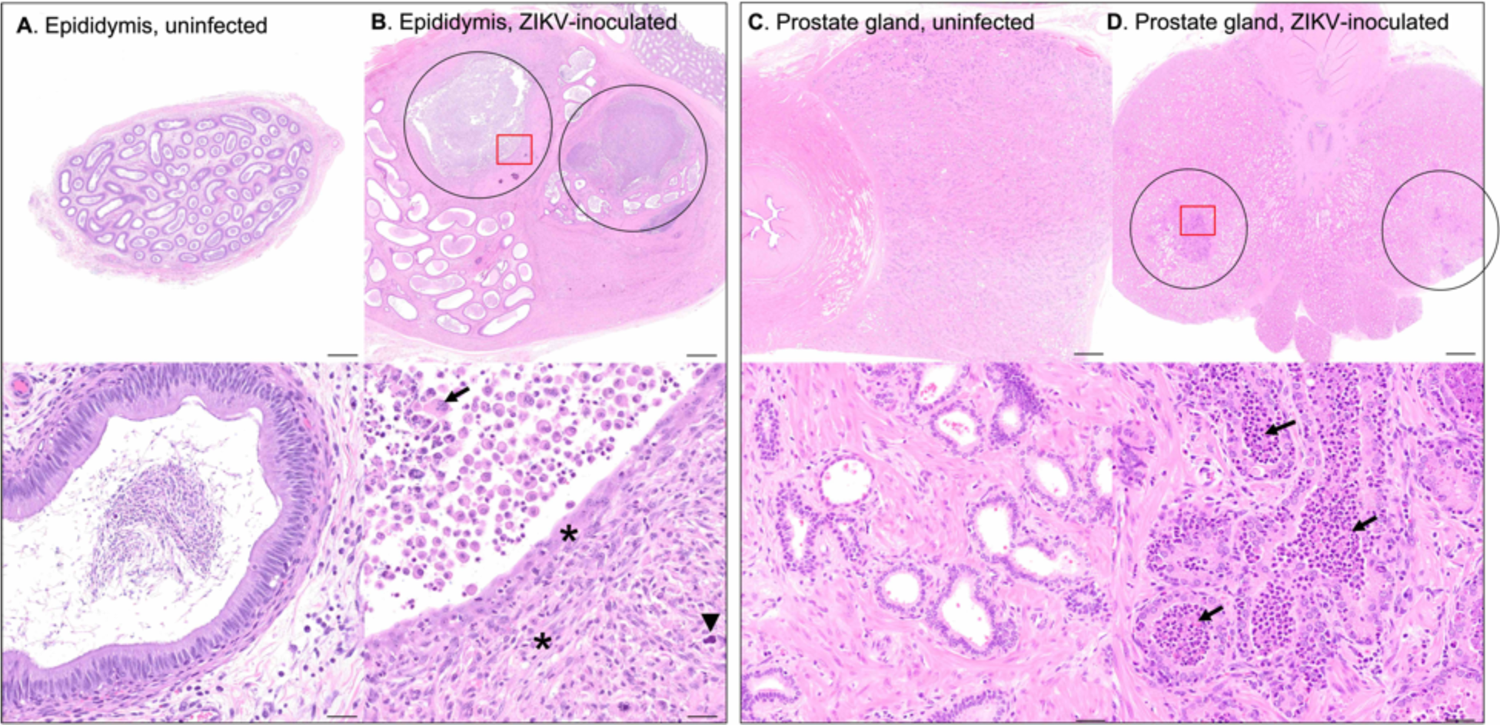
Photomicrographs of H&E-stained genital tissues from uninfected & ZIKV-inoculated male rhesus macaques A.) Normal epididymis from an uninfected control animal. **B.)** Epididymis from a ZIKV-inoculated rhesus macaque with duct rupture and pyogranulomatous epididymitis (circled). Remaining epididymal ducts are dilated and tortuous with sperm stasis. In the 20X image (lower panel, area denoted by red box in upper panel) normal duct epithelial architecture is lost with replacement by fibroplasia (asterisks) and mineralization (arrowhead) with neutrophils, multinucleated giant cells (arrow), and necrotic debris. This correlates with virology data, where macaques with detectable epididymal ZIKV RNA exhibited higher histology scores than those without detectable virus (data not shown, linear model, p = 0.008). **C.)** Normal prostate gland from an uninfected control animal. **D.)** Prostate gland from a ZIKV-inoculated macaque with mild to moderate prostatitis (circled). In the 20X image (lower panel, area denoted by red box in upper panel) glandular lumens are expanded or replaced by sloughed cells, necrotic debris, neutrophils, and macrophages (arrows). Scale bars: upper images 1mm; lower images 20μm.

The most common histologic finding in the prostate gland was mild to moderate prostatic inflammation (4/28 animals) characterized by periglandular aggregates of lymphocytes, macrophages, and scattered neutrophils. Affected glands were often expanded by sloughed cells, necrotic debris, neutrophils, and macrophages (**Figure 6C, D**). While both ZIKV RNA and infectious virus were frequently detected within the seminal vesicle, no significant microscopic lesions were noted in this tissue in ZIKV-inoculated animals. There was variably severe mineralization of secretory product; however, this was also present in control tissues and is a very common, clinically insignificant background lesion in the seminal vesicle of sexually mature macaques (25).

Microscopic evaluation of macaque testes was complicated by the use of conventional formalin fixation (where special fixatives are preferred) (26), lack of serial sectioning, small sample size, poor preservation and/or crush artifact in some samples. Furthermore, rhesus macaques, in contrast to cynomolgus macaques, are seasonal breeders (26), which results in reduced spermatogenesis and low hormone levels out of season (27). As a result of these complicating factors, only significant testicular lesions, including inflammation, necrosis, or evidence of sperm stasis (as evidenced by luminal aggregation of spermatozoa within seminiferous tubules or rete testes and/or luminal macrophages with engulfed spermatozoa) were scored for statistical analyses. Using this criteria, testicular lesions were uncommon and mild in ZIKV-inoculated animals.

In summary, we identified a significant association between ZIKV-infected, sexually mature male macaques and pathologic microscopic lesions in the epididymis and prostate gland, predominantly between 1 and 20 DPI, that were absent in non-infected animals.

## Discussion

In this study, we investigated a poorly understood aspect of human ZIKV infection. We evaluated role of the male reproductive tract in the context of viral persistence in genital tissues, potential for sexual transmission, and microscopic lesions along with their potential effects on fertility. This study further emphasizes the utility and relevance of macaque models of ZIKV infection, as we were able to evaluate archived reproductive tissue samples from more than 50 male macaques, which are generally very difficult to obtain from ZIKV-infected men. Intravenous or subcutaneous inoculation of Brazilian or Puerto Rican ZIKV of male rhesus and cynomolgus macaques produced asymptomatic infection with viremia lasting from 4 to 14 DPI and peaking at 4 to 5 DPI. These findings are consistent with those from adult, non-pregnant ZIKV-infected humans (28, 29), as well as published data from NHP models of ZIKV infection (13–15,18,30–32).

Our findings, including the detection of both ZIKV RNA and infectious virus in male macaque genital tissues, and ZIKV RNA (but not infectious virus) in the two available semen samples, support the hypothesis that male reproductive tract serves as a reservoir for ZIKV (33). Our findings are also consistent with published data in NHP, which, while sparse, have demonstrated ZIKV RNA in macaque semen for up to 28 DPI (30), and in the testis (17, 30), seminal vesicle (18, 30), and prostate gland (18,30,34) from 4 to 35 DPI, frequently after the resolution of viremia. We detected ZIKV RNA most frequently in the epididymis and seminal vesicle, with the highest absolute magnitude occurring in the epididymis. A higher peak ZIKV viremia, larger viremia AUC, and detection of ZIKV RNA in the epididymis and seminal vesicle correlated with sexual maturity in macaques. Infectious ZIKV was cultured from at least one genital tissue, most frequently in the seminal vesicle and epididymis, in 38% (18/48) of ZIKV-inoculated macaques. ZIKV RNA persisted in male macaque genital tissues for up to 60 DPI, about 6 weeks after the resolution of viremia at 14 DPI in most animals in this study. Our results extend knowledge on the duration of persistence since 35 DPI was the longest previously documented duration of ZIKV RNA in male macaque genital tissues (18). As 60 DPI was the latest time point assessed in this study, it is possible that persistence of ZIKV RNA in male macaque genital tissues may be even more prolonged. This finding suggests that the potential for sexual transmission of ZIKV remains even after viremia has resolved, which raises significant concerns regarding the risks of male sexual transmission to both men and periconceptional and non-pregnant women, as well as in the context of assisted fertility procedures such as sperm donation. With the exception of one reported case of congenital Zika syndrome arising from sexual transmission of ZIKV from an infected man to his naïve, pregnant wife (23), the relationship between sexual and vertical transmission from mother to fetus is poorly understood, and further study is needed.

Semen obtained from symptomatic convalescent men can harbor both ZIKV RNA and infectious virus after the resolution of viremia (35–37). The latest documented report of human sexual transmission was 44 days after the onset of symptoms (38). One study evaluating ZIKV infection in men reported that older age, infrequent ejaculation, and the presence of certain symptoms (i.e., conjunctivitis) at the time of initial illness were associated with prolonged sexual shedding of ZIKV RNA (37). Additional data suggest that persistent ZIKV infection of the male reproductive tract may stimulate a prolonged immune response. Long-term male shedders, defined as men with detectable ZIKV RNA in their semen for greater than 3 months, had significantly higher seminal leukocyte counts and pro-inflammatory cytokines including IL-6 and IL-8 when compared to short-term shedders (39). It is unclear whether the absence of infectious virus from semen samples in the present study denotes a lack of transmission potential, prior infection followed by clearance below the limit of detection, or a methodological artifact of infectious virus decay after a freeze-thaw cycle that would result in a false negative in a sample of low titer. Unfortunately, metadata (including age, ZIKV strain/dose, route of inoculation, and duration of infection) were unavailable for the animals from which these samples were collected, so it is possible that viral migration to the male reproductive tract occurred early during infection, and by the time of necropsy, when a “snapshot” is taken, ZIKV RNA or infectious virus levels declined to low or undetectable levels.

The male ejaculate is composed of both cellular and fluid components. The cellular component comprises spermatozoa, and white blood, desquamated germ, and epithelial cells. The fluid component contains secretions from accessory sex glands, primarily the seminal vesicle and prostate gland, and to a lesser extent, the bulbourethral gland and epididymis (**Figure 7**). Either or both components could harbor infectious ZIKV and contribute to sexual transmission. Published data regarding specific cellular tropisms of ZIKV are somewhat conflicting, as detailed below. *In vitro* studies using human cells have variously demonstrated that primary testicular germ cells (40–42), Sertoli cells (41–45), peritubular myoid cells (42), epididymal epithelial cells (46), fibroblasts (41), and epithelial and mesenchymal stem cells of the prostate gland (47) are susceptible to ZIKV infection. Following ZIKV inoculation of *ex vivo* testicular explants, macrophages, and peritubular cells are most frequently infected, with fewer Leydig and Sertoli cells infected (33). ZIKV inoculation of human testicular organoids (HTO) results in productive ZIKV infection, with decreased HTO survival and reduced expression of spermatogonial, Sertoli, and Leydig cell markers (48). *In vivo* studies using immunodeficient mice have similarly demonstrated virus localization to sperm precursors (19), Sertoli cells (19, 49), interstitial Leydig cells (19, 21), epididymal epithelium (21,49–52), prostatic epithelium (34), and the cell-free seminal plasma fraction of the murine ejaculate (49), from 3 to 33 DPI (50–52).

**Figure 7.**
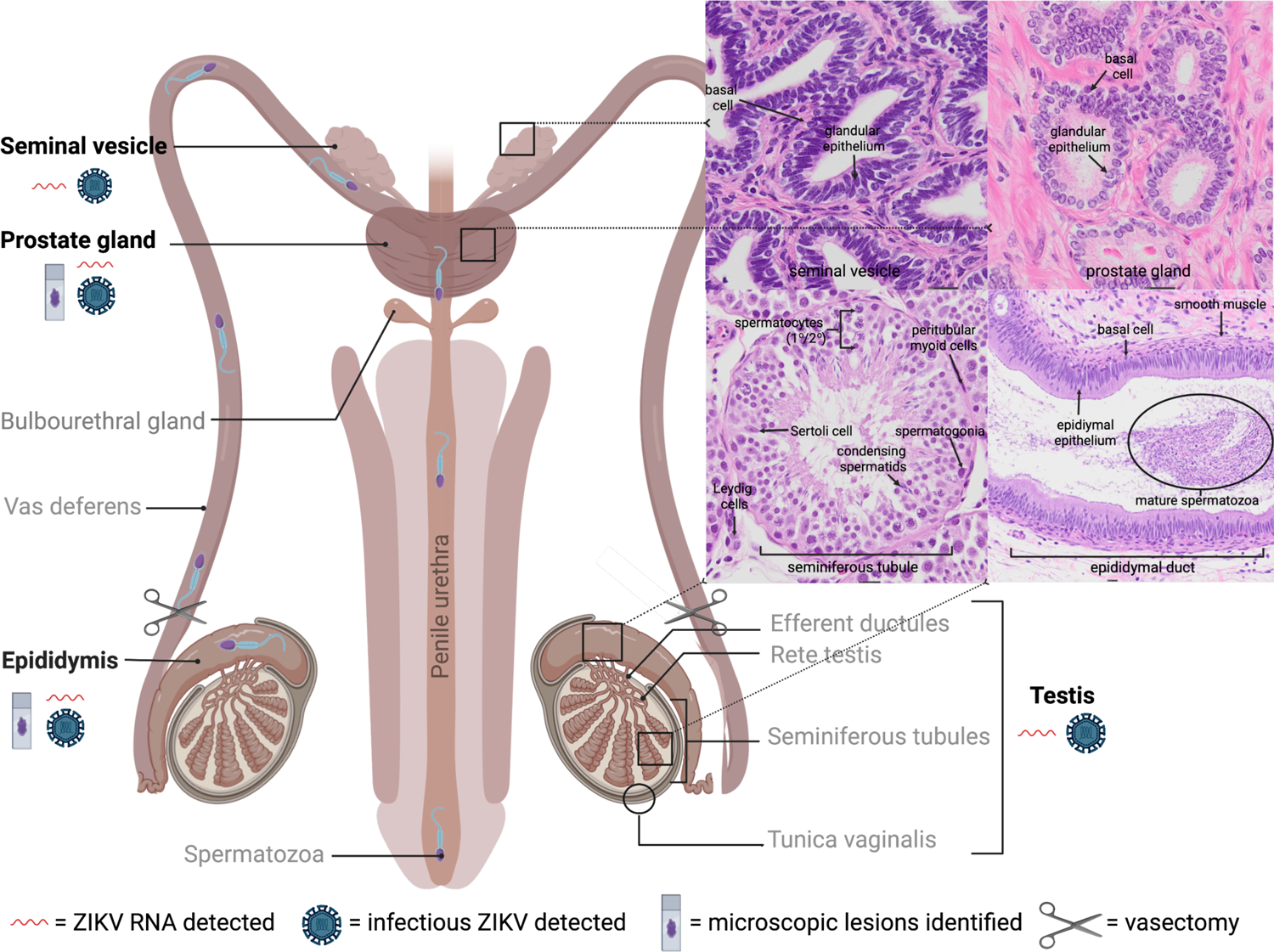
Normal anatomy and histology of the sexually mature male primate reproductive tract with summary of results. Tissues that were sampled are in bold. Insets (denoted by black boxes on diagram) depict normal microscopic features of each tissue. Mature spermatozoa develop in stages from germ cells (spermatogonia) in testicular seminiferous tubules (supported by Sertoli cells), undergo maturation and storage in the ducts of the epididymis, and travel via the vas deferens through the accessory sex glands and into the urethra for ejaculation (74, 75). Both ZIKV RNA and infectious virus were detected in the testis, epididymis, seminal vesicle, and prostate gland. Microscopic lesions were noted in the sexually mature epididymis and prostate gland. Created with BioRender.com.

Viral antigen was identified in mature spermatozoa from the semen sample of a ZIKV-infected man (53), and ZIKV shedding in human semen has been shown to correspond with the duration required for the human spermatogonial life cycle, which is approximately 74 days (54). Infectious virus and ZIKV RNA have been isolated from the semen of vasectomized men for up to 69 and 96 days after symptom onset, respectively (55, 56) ZIKV RNA and infectious virus levels are significantly reduced in vasectomized versus non-vasectomized ZIKV-infected men (37). The single vasectomized rhesus macaque in the present study had detectable ZIKV RNA in all 4 genital tissues, and infectious virus was cultured from the seminal vesicle (semen was not available from this animal). A vasectomy is a surgical sterilization procedure that entails cutting the vas deferens to prevent sperm from leaving the epididymis and entering the male ejaculate (57). Detection of ZIKV within semen and/or accessory sex glands of vasectomized males suggests that infected spermatozoa are not required for sexual transmission and that infectious cells/fluids from the epididymis, seminal vesicle and/or prostate gland may also play a significant role in sexual transmission. Furthermore, mature spermatozoa lack endoplasmic reticula, Golgi apparati, and tRNAs and are considered transcriptionally inactive (58) and therefore could not be expected to be infected with and produce ZIKV.

Results from *in vitro* and *in vivo* experiments as well as observations in humans suggest that chronic ZIKV infection in males could result in sexual transmission of virus; however, while germ cells and later sperm precursors harbor ZIKV, mature spermatozoa are probably not the only source of infectious ZIKV in semen or the sole contributor to sexual transmission. Rather, epididymal epithelial cells, leukocytes like macrophages, and other transcriptionally active cells are more probable sources of replicating ZIKV in semen and likely play a crucial role in sexual transmission of ZIKV (51). Our results support this conclusion. Using ISH we showed that, in addition to testicular germ cells and 1°/2° spermatocytes, ZIKV RNA localized to macaque Sertoli cells (which are modified epithelial cells), epididymal epithelial cells, and glandular epithelial cells within the seminal vesicle and prostate gland during both acute and chronic stages of disease. Taken together, our findings and published data suggest that ZIKV is capable of breaching blood-testis and/or blood-epididymal-barriers to replicate in multiple cell types and persist in the male reproductive tract, though specific mechanisms by which ZIKV enters these cells and establishes persistence remain undefined. Additionally, our observation that ZIKV RNA localized most frequently to testicular stem-like cells including germ cells and spermatocytes is in agreement with previous macaque studies demonstrating an apparent viral preference for stem cells, such as fetal neural precursor cells in macaque fetuses exposed to ZIKV prenatally (59).

ZIKV is clearly gonadotropic in men, but information regarding genitourinary sequelae and effects on fertility are lacking in both humans and animal models of human ZIKV infection. Hematospermia, prostatitis, painful ejaculation, penile discharge, dysuria, low sperm counts and sperm motility issues are occasionally reported in ZIKV-infected men (8,9,60,61); however, human data remain sparse, as obtaining genital biopsy specimens from men is invasive and not typically performed without a significant medical reason (62). Up to 80% of ZIKV-infected humans remain asymptomatic, and gauging the true prevalence of ZIKV in semen, understanding associated viral kinetics, recognizing factors influencing male sexual transmission, characterizing male genitourinary lesions, and identifying potential effects on fertility and risk factors associated with sperm donation and assisted fertility procedures have proven quite difficult (8).

ZIKV infection of immune suppressed mice induces testicular and epididymal damage progressing to atrophy, with microscopic evidence of orchitis and epididymitis, low serum testosterone, and decreased fertility (19,21,40,44,46,50–52,63). The prostate gland and seminal vesicles are typically spared (51, 52), though ZIKV-associated prostatitis is occasionally identified both immunodeficient mice and immunocompetent macaques (34). In contrast to mice, genitourinary lesions and infertility are usually not major features in macaques inoculated with ZIKV (17,18,30). This discrepancy may have to do with the use of immunodeficient knockout mice versus immune competent macaques and/or the low sample sizes typically used in NHP studies, which often do not provide sufficient statistical power to identify infertility if rare.

In the present study, microscopic lesions were relatively uncommon in the reproductive tract of ZIKV-inoculated male macaques; however, microscopic lesion severity scores for both the epididymis and prostate gland were significantly higher in sexually mature ZIKV-inoculated male macaques versus uninfected, age-matched controls and ZIKV-inoculated, sexually immature animals. Specifically, ZIKV-inoculated sexually mature macaques exhibited sporadic, variably severe epididymitis and/or prostatic inflammation. Statistically, the macaques with detectable ZIKV RNA in the epididymis exhibited significantly higher histology scores than those without detectable virus. These findings further underscore the potential importance of the epididymis in the context of ZIKV persistence and shedding, sexual transmission, and associated genitourinary pathology.

The moderate to severe epididymal inflammation noted in occasionally within these ZIKV-inoculated macaques is suggestive of epithelial damage that may have been ZIKV-induced, followed by epididymal duct rupture and exposure of luminal contents including sperm to the surrounding tissue, which incites a severe inflammatory response. Epididymitis in primates has several documented etiologies, including ascending bacterial or fungal infection from sexually transmitted or urinary tract infections, trauma/obstruction and, less commonly, viral infection as caused by orthorubulaviruses (mumps), adenoviruses, or enteroviruses (64). Prostatic periglandular and/or perivascular lymphocytic infiltrates are common, clinically insignificant background findings in macaques, however, the presence of neutrophilic inflammation and necrotic debris, as seen here, are not (25). As noted for epididymitis, ascending bacterial infection associated with recurrent urinary tract infection is the most frequently reported cause of prostatitis in humans (65). The same is likely also true in macaques, although prostatic inflammation is much less common in NHP (66). Although ascending bacterial infection is a more commonly reported cause of both epididymitis and prostatitis than viral infection (64), the macaques in this study lived controlled environments, underwent regular physical examinations and blood work, and exhibited no evidence of trauma or infection (other than ZIKV) at necropsy or by microscopic evaluation. While we cannot definitively prove causation, there is an association between ZIKV and histologic genital lesions which warrants further study. Furthermore, epididymitis is associated clinically with male infertility (64), and chronic prostatitis in humans is a well-known precursor to prostatic carcinoma (67), so a developing a thorough understanding of potentially ZIKV-induced pathologic lesions can inform long-term implications for the health of men in ZIKV-endemic regions. Further study, including detailed analysis of semen from ZIKV infected macaques and humans, is necessary.

Sexual maturity also had a significant effect on our results. Sexually mature male macaques were more likely to harbor persistent ZIKV in the reproductive tract, particularly in the epididymis or seminal vesicle. Furthermore, significant histopathologic lesions only occurred in sexually mature male macaques inoculated with ZIKV. The explanation for this observation likely relates to anatomic and physiologic differences between sexually immature and mature males, such as the types and relative differentiation of cells present in genital tissues and accessory sex glands, receptor expression, and/or hormone production. In contrast to the post-pubertal macaque testis, sexually immature seminiferous tubules possess smaller diameter lumens and contain only Sertoli cells and undifferentiated spermatogonia (11). Epididymal ducts are similarly reduced in diameter, lined by small, flattened epithelial cells, and surrounded by increased fibrous connective tissue (11). Glands of the sexually immature seminal vesicle and prostate gland are also narrowed, lack luminal secretions, and are lined by low cuboidal to flattened epithelial cells (11).

In conclusion, we show here that the male macaque reproductive tract serves as a reservoir for ZIKV RNA and infectious virus, that epithelial cells and mesenchymal/stem cells of the testis, epididymis, seminal vesicle, and prostate gland can harbor ZIKV RNA, and that ZIKV infection is associated with microscopic lesions in the epididymis and prostate gland. The immune-privileged, inherently immunosuppressive nature of the testes and epididymis (68) likely promotes ZIKV persistence and sexual transmission of infectious virus beyond the acute stage of infection. Overall, sexually mature males are at significantly higher risk for genital ZIKV persistence and urogenital sequelae, though mechanisms of viral entry into the male reproductive tract and the pathogenesis of injury to genital tissues remain unclear. Taken together, our results support the hypotheses that 1) the male genital tissues including accessory sex glands such as the seminal vesicle and prostate gland serve as a reservoir and probable replication site for ZIKV, and 2) that genital lesions and impaired male fertility are possible, if not likely, sequelae to ZIKV infection. Furthermore, our identification of ZIKV RNA in frozen semen samples, and detection of ZIKV RNA and infectious virus in frozen genital tissue samples for up to 60 DPI and 50 DPI respectively, indicate that freezing is not a viable method of destroying ZIKV. This has significant implications for the safety of assisted fertility procedures involving donated reproductive tissues such as sperm, oocytes and embryonic tissue, where ZIKV screening and testing is recommended but not required by the U.S. Food and Drug Administration and is performed at the discretion of individual clinics (22, 69).

By extrapolating our findings from ZIKV-infected macaques, we can significantly increase our understanding of persistent ZIKV infection in men. This was an opportunistic study where we leveraged archived macaque tissues originating from different studies where the animals were exposed to variable experimental conditions. Additional, controlled experiments are clearly needed, particularly to define: 1) mechanisms of viral entry into the male reproductive tract; 2) effects of ZIKV infection on the histomorphology of genital tissues and fertility, including detailed analysis of semen, in sexually mature ZIKV-infected males; and 3) the relationship between sexual transmission to naïve periconceptional woman and risks of vertical transmission to the fetus.

## Materials and Methods

### Study Design

The tissues evaluated in this study were from healthy male rhesus (*Macaca mulatta*) and cynomolgus (*Macaca fascicularis*) macaques, born at their respective National Primate Research Centers (**Table 1**). All studies were approved by the appropriate Institutional Animal Care and Use Committees (IACUC). Archived reproductive tissues, including testes, epididymis, seminal vesicle, and prostate gland, from 51 ZIKV-infected and 8 age-matched, uninfected, male rhesus and cynomolgus macaques from past or ongoing collaborative research projects were kindly donated by the California (N = 35) (13,31,32), Tulane (N = 10) (70), Wisconsin (N = 6) (71, 72), and Washington (N = 1 uninfected control rhesus macaque) NPRCs. These animals, aged 2 to 15 years old, were inoculated IV or SC with variable ZIKV strains and doses, humanely euthanized, and necropsied at 1 to 60 DPI. When possible, testis, epididymis, seminal vesicle, and prostate gland were examined; however, a full complement of genital tissues was not collected from each animal. Among rhesus macaques from the CNPRC, 9 animals that failed to become viremic upon initial inoculation were reinoculated, and 4 animals were immune suppressed via CD8+ T-cell depletion 4 weeks after inoculation to evaluate the potential for viremia resurgence, which did not occur. Plasma/serum and or viremia data was available from only a subset of these animals (N = 36). Uninfected, age-matched control tissue was obtained from colony management culls at the California and Washington NPRCs.

### Necropsy, tissue collection and histopathology

All necropsies were performed by a board-certified veterinary pathologist and 1 to 2 technicians. Macaques were euthanized with an overdose of sodium pentobarbital. The veterinary pathologist evaluated each tissue *in situ* prior to excision. Technicians trimmed each tissue using separate forceps and razor blades to minimize risks for cross-contamination. Male reproductive tissues, including testis, epididymis, seminal vesicle, and prostate gland were collected. Tissues were collected for viral analyses in RNAlater (Thermo Fisher Scientific, Waltham, MA) according to the manufacturer’s instructions. Extra available samples were snap-frozen and stored at −70°C. Tissues for histopathology were preserved in 10% neutral-buffered formalin (Thermo Fisher Scientific), paraffin-embedded, thin sectioned (5 μm), routinely stained with hematoxylin and eosin (H&E) and evaluated by a board certified veterinary anatomic pathologist, generating a cumulative abnormality score from 0 (normal) to 5 (markedly abnormal) (**Supplemental Table 3)**.

### Isolation and quantification of viral RNA from plasma and male genital tissues

ZIKV RNA was isolated from samples and measured in triplicate by qRT-PCR according to methods described previously (13). Briefly, EDTA-anticoagulated whole blood was centrifuged for 10 minutes at 800 g and the resulting plasma fraction was stored at −70°C. RNA was extracted from plasma according to the manufacturer’s instructions using the MagMAX Express-96 Deep Well Magnetic Particle Processor (Thermo Fisher Scientific). Solid tissues frozen in RNAlater were thawed and homogenized to a liquid state using a 5 mm steel ball, Qiazol lysis reagent and the Qiagen/Retsch TissueLyser II (all from Qiagen, Germantown, MD). RNA was extracted from homogenized tissue supernatants using the viral RNA universal mini kit (Qiagen) or the automated QIAcube (Qiagen). All RNA extracts were eluted in 60 μL of diethyl pyrocarbonate (DEPC)-treated water for storage at −70°C prior to quantification and were tested in triplicate using an Applied Biosystems ViiA 7 RT-qPCR machine (Thermo Fisher Scientific). Viral RNA levels were calculated in RNA copies by comparing the average of each triplicate from a sample to the standard curve generated with each PCR plate. Levels of ZIKV RNA in samples are expressed as mean log_10_ RNA copies per mL fluid or mg tissue. The limit of detection (LOD) varied depending on the volume/weight of tissue sampled and volume of Qiazol needed to homogenize to liquefaction, with means of 1.9 log_10_ RNA copies/mL and 2.8 log_10_ RNA copies/mg tissue. A sample was considered positive when 2 of 3 or all 3 replicates yielded an RNA copy value. When a sample exhibited an inconsistent qRT-PCR signal (1 of 3 replicates positive) retesting was performed, generally on a different aliquot, if available. If the retest result was negative (3 of 3 replicates), the sample was considered negative. If 1 of 3 replicates remained positive but was within 1 log_10_ RNA copies/mg tissue or mL fluid of the LOD, the sample was also considered negative.

### Infectious Zika virus quantification by plaque assay

Infectious ZIKV was detected using a Vero cell (American Type Culture Collection, Manassas, VA) plaque assay, as described previously (13). Briefly, confluent 12-well Vero plates were inoculated with 250 μL of 1:10 and 1:20 dilutions of macaque tissue homogenate in Dulbecco’s Modified Eagle Medium (DMEM) (Thermo Fisher Scientific) supplemented with 2% fetal bovine serum and allowed to absorb at 37°C for 1 hour. After incubation, each cell monolayer was overlaid with 1 ml 0.4% agarose (liquefied 10% agar, ultrapure agarose [Invitrogen, Carlsbad, CA] diluted in 42°C DMEM) and allowed to solidify. The plates were incubated at 37°C for 7 days. Cell monolayers were then fixed with 4% formalin for 30 minutes, agar plugs were gently removed, and viable cells were stained with 0.05% crystal violet (Sigma, St. Louis, MO) in 20% ethanol. Viral titers were recorded as the reciprocal of the highest dilution where plaques are noted. The limit of detection of the assay was 0.4 PFU/mg tissue. Plaque assays were only performed on tissues that tested positive via qRT-PCR and were not performed on samples that contained less than 3 log_10_ genomes/mg tissue since our previous work showed those samples were not likely to contain infectious ZIKV. Two replicate titrations were performed for each sample and the replicate measurements were averaged.

### Viral RNA detection by *in-situ* hybridization (ISH)

Colorimetric *in-situ* hybridization (ISH) was performed manually on superfrost plus slides (Thomas Scientific, Swedesboro, NJ), according to the manufacturer’s instructions (73), using the RNAscope 2.5 HD Red Reagent Kit (Advanced Cell Diagnostics, Newark, CA) and RNAscope Probe V-ZIKVsph2015 (Advanced Cell Diagnostics). Briefly, each 5 μm section of formalin-fixed, paraffin embedded tissue was pretreated with heat and protease, followed by ZIKV probe (GenBank accession number KU321639.1 [complete genome]; 70 pairs; target region 130-4186) hybridization for 2 hours at 40°C, a cascade of signal amplification molecules, and signal detection. Slides were counterstained with hematoxylin and mounted with xylene based EcoMount (BioCare Medical, Pacheco, CA). A probe designed to detect bacterial dapB (Advanced Cell Diagnostics, Newark, CA) was used as a negative control.

A section of spleen from a ZIKV-infected rhesus macaque euthanized 4 DPI was used as a positive control. ISH was only performed on tissues that tested positive via qRT-PCR and was not performed on samples that contained less than 5 log_10_ genomes/mg tissue since previous (unpublished) work showed those samples are unlikely to show detectable signal. Positive staining was identified as red cytoplasmic or perinuclear staining. When possible, ISH-positive cell types were identified by the pathologist based on tissue location and cell morphology.

### Data analyses

A noteworthy feature of this project is that it leverages existing, archived tissues, including tissues from various studies performed at three other primate centers in addition to our CNPRC (Tulane, Wisconsin and Washington NPRCs). While no additional animals were infected to conduct this project, samples originated from macaques exposed to differing experimental conditions, including frequency and route of inoculation, viral strain and dose, primate species, and immune-suppression via CD8+ T-cell depletion (**Table 1**). To account for these differing experimental conditions, we investigated the association between the variables with tests for independence. We conducted a univariate analysis using Wilcoxon rank test for continuous variables, fisher’s exact test for categorical, and linear regression for the histology scores. All the statistical analysis were performed in R-studio. Graphs were created using GraphPad Prism. P-values of less than or equal to 0.05 were considered statistically significant.

## Acknowledgements

We thank K. Jackson for assistance completing *in-situ* hybridization; J. Vidal for assistance with histopathology interpretation; J. Go and the Washington National Primate Research Center Tissue Distribution Program, and the pathology staff at the UC Davis School of Veterinary Medicine and the California National Primate Research Center.

## Financial Support

This work was supported by the Office of Research Infrastructure Program, Office of the Director, National Institutes of Health Award Numbers P51OD011107 (CNPRC), P51OD011106 (WNPRC), P51OD011104 (TNPRC), and R01AI116382 (David H. O’Connor); the University of California, Davis Graduate Student Support Program; the National Institutes of Health Comparative Medical Science Training Program Award Number T32 OD 011147; and the U.S. Army Medical Center of Excellence Long Term Health Education and Training Program.

## Disclosures

The authors do not declare any conflict of interest. Funding sources did not influence experimental design and analysis/interpretation of results or impact the decision to publish.

## Data Availability

All data contributing to the generation of figures and analyses described herein are available upon request from the corresponding author.

## Author Contributions

Conceptualization: EEB, LLC, KVR, PP

Data curation: EEB, AS, DD, NM, BS, AP, DO, MB, MKK, JPG

Formal analysis: EEB, JPG

Funding acquisition: EEB, LLC, KVR Investigation: EEB, LLC, KVR, PP

Methodology: EEB, LLC, KVR, PP, AS, DD, NM, BS, AP, DO, MB, MKK, JPG

Project administration: EEB, LLC, KVR Supervision: EEB, LLC, KVR, PP Writing – original draft: EEB

Writing – review & editing: EEB, LLC, KVR, PP, AS, DD, NM, BS, AP, DO, MB, MKK, JPG

## Notes

### Competing Interest Statement

The authors have declared no competing interest.

