## Supplemental Tables for "Zika virus persistence in the male macaque reproductive tract"

**Supplemental Table 1:** **Experimental conditions exerting a significant effect on dependent variables in ZIKV inoculated male macaques**. All statistical tests performed using R. Only significant and borderline P-values reported here. P value ≤ 0.05 is significant, P value 0.05-0.08 is borderline. CI is confidence interval; Ab is antibody; DPI is days post-inoculation; AUC is area under the curve. ^1^Intravenous (IV) as reference.

| Dependent Variable | Experimental Condition | Test | Estimate  (95%CI) | P-value |
| --- | --- | --- | --- | --- |
| Viremia positive/negative | Administration of anti-ZIKV Ab | Fisher’s exact | 0  (0.0, 1.19) | 0.03 |
| Peak viremia magnitude | Administration of anti-ZIKV Ab | t-test | 2.03  (0.93, 3.07) | 0.02 |
| Peak viremia (DPI) | Administration of anti-ZIKV Ab | t-test | -9.23  (-20.90, -11.49) | 0.0005 |
| Overall viremia duration | Administration of anti-ZIKV Ab | t-test | -7.69  (-15.86, -8.35) | 0.0001 |
| Peak viremia magnitude | ZIKV Dose | Kruskal Wallis | 19.73 (chi square) | 0.03 |
| Peak viremia magnitude | Route of inoculation^1^ | t-test | 2.62  (0.21, 1.89) | 0.02 |
| Viremia AUC | Reinoculation | t-test | 3.89  (10.68, 43.94) | 0.002 |
| Detection of ZIKV RNA in the male reproductive tract | Route of inoculation^1^ | Fisher’s exact | 8.71  (1.52, 94.17) | 0.01 |
| Detection of ZIKV RNA in epididymis | Route of inoculation^1^ | Fisher’s exact | 13.5  (2.23, 115.66) | 0.001 |
| Detection of ZIKV RNA in epididymis | Administration of anti-ZIKV Ab | Fisher’s exact | 0.073  (0.001, 0.73) | 0.01 |
| Detection of ZIKV RNA in seminal vesicle | Route of inoculation^1^ | Fisher’s exact | 6.11  (1.41, 31.35) | 0.01 |
| Histology score, epididymis | Route of inoculation^1^ | Simple logistic regression | 1.03  (0.36, 1.70) | 0.004 |
| Histology score, prostate gland | Route of inoculation^1^ | Simple logistic regression | 0.56  (0.01, 1.10) | 0.05 |
| Histology score, epididymis | Reinoculation | Simple logistic regression | 1.62  (0.88, 2.36) | 0.002 |
| Histology score, prostate gland | Reinoculation | Simple logistic regression | 0.95  (0.30, 1.61) | 0.01 |
| Histology score, prostate gland | Administration of anti-ZIKV Ab | Simple logistic regression | -1.32  (-1.96, -0.68) | <0.0001 |

**Supplemental Table 2:** **Table summarizing statistically significant variables in ZIKV inoculated male macaques.** Statistical tests performed using R-studio or Graphpad Prism. Only significant P-values reported here. P value ≤ 0.05 is significant, P value of 0.05-0.08 is borderline; CI is confidence interval; DPI is days post-inoculation; AUC is area under the curve.

| Dependent variable | Independent variable | Test | ESTIMATE  (95% CI) | P-value |
| --- | --- | --- | --- | --- |
| Peak viremia magnitude | Sexual maturity | Ordered logistic regression model | 1.19  (0.46, 1.93) | 0.002 |
| Viremia AUC | Sexual maturity | Ordered logistic regression model | -21.62  (-41.2, -2.03) | 0.03 |
| Detection of ZIKV RNA in epididymis | Magnitude peak viremia | Ordered logistic regression model | 0.24  (0.11, 0.37) | 0.0007 |
| Detection of ZIKV RNA in male reproductive tract | Sexual maturity | Ordered logistic regression model | 0.33  (0.06, 0.61) | 0.02 |
| Detection of ZIKV RNA in epididymis | Sexual maturity | Ordered logistic regression model | 0.71  (0.46, 0.95) | <0.0001 |
| Detection of ZIKV RNA in seminal vesicle | Sexual maturity | Ordered logistic regression model | 0.59  (0.28, 0.88) | 0.0005 |
| Detection of ZIKV RNA in seminal vesicle | Days post-inoculation | Ordered logistic regression model | -0.02  (-0.02, -0.006) | 0.005 |
| Detection of ZIKV RNA in seminal vesicle from 1-20 DPI | Detection of ZIKV RNA in seminal vesicle from 21-40 DPI | Mann-Whitney | - | 0.02 |
| Detection of ZIKV RNA in seminal vesicle from 1-20 DPI | Detection of ZIKV RNA in seminal vesicle from 41-60 DPI | Mann-Whitney | - | 0.03 |
| Histology score epididymis (ZIKV-inoculated) | Histology score epididymis (uninfected controls) | Mann-Whitney | - | 0.02 |
| Histology score prostate (ZIKV-inoculated) | Histology score prostate (uninfected controls) | Mann-Whitney | - | <0.0001 |
| Histology score epididymis | Sexual maturity | Linear model | 0.92  (0.15, 1.68) | 0.02 |
| Histology score prostate | Sexual maturity | Linear model | 1.17  (0.62, 1.71) | 0.0001 |
| Histology score epididymis | Detection of ZIKV RNA in epididymis | Linear model | 1.09  (0.34, 1.84) | 0.01 |
| Histology score epididymis | Days post-inoculation | Linear model | -0.03  (-0.05, -0.007) | 0.02 |
| Histology score prostate | Days post-inoculation | Linear model | -0.02  (-0.05, -0.003) | 0.03 |

**Supplemental Table 3**: Histologic scoring criteria for the male macaque genital tissues

|  | **Testis** | **Epididymis** | **Seminal vesicle** | **Prostate gland** |
| --- | --- | --- | --- | --- |
| **0** (none) | No significant lesions | No significant lesions | No significant lesions | No significant lesions |
| **1** (minimal) | Lesions minimal and significance questionable (affecting <5% of the visible surface area): Rare perivascular/ peritubular mononuclear infiltrates (less than 3 small foci); and/or mild evidence of sperm stasis (rete testes or efferent ducts exhibit sperm aggregation with debris, macrophages, and multinucleated giant cells +/- engulfed sperm) | Lesions minimal and significance questionable (affecting <5% of the visible surface area): Rare perivascular/ periductular mononuclear infiltrates (less than 3 small foci); and/or mild evidence of sperm stasis (dilated epididymal ducts lacking sperm with replacement by debris, macrophages, and multinucleated giant cells +/- engulfed sperm) | Lesions minimal and significance questionable: Rare perivascular/ peritubular mononuclear infiltrates (less than 3 small foci) | Lesions minimal and significance questionable (affecting <5% of the visible surface area): Rare perivascular/periglandular mononuclear infiltrates (less than 3 small foci); and/or foci of increased fibrous connective tissue |
| **2** (mild) | Mild lesions not observed in control animals (affecting 5-10% of the visible surface area): As above with more frequent perivascular/peritubular mononuclear infiltrates; and/or mild mixed inflammation, hemorrhage/edema; and/or mild evidence of seminiferous tubule degeneration | Mild lesions not observed in control animals (affecting 5-10% of the visible surface area): As above with more frequent perivascular/periductular mononuclear infiltrates; and/or mild mixed inflammation, hemorrhage/edema; and/or mild evidence of ductular epithelial degeneration | Mild lesions not observed in control animals (affecting 5-10% of the visible surface area): As above with more frequent perivascular/peritubular mononuclear infiltrates; and/or mild mixed inflammation, hemorrhage/edema | Mild lesions not observed in control animals (affecting 5-10% of the visible surface area): As above with more frequent perivascular/ periglandular mononuclear infiltrates; and/or mild mixed inflammation, hemorrhage/edema; and/or expansion of glandular lumens by necrotic debris, neutrophils, and mononuclear inflammatory cells; and/or occasional foci of mineralization |
| **3** (moderate) | Moderate lesions not observed in control animals (affecting 10-20% of the visible surface area): As above with multiple larger foci of mixed inflammation; and/or rare, small granulomas; and/or scattered foci of mineralization; and/or seminiferous tubule necrosis | Moderate lesions not observed in control animals (affecting 10-20% of the visible surface area): As above with multiple larger foci of mixed inflammation; and/or rare, small granulomas; and/or scattered foci of mineralization; and/or ductular epithelial necrosis | Moderate lesions not observed in control animals (affecting 10-20% of the visible surface area): As above with multiple larger foci of mixed inflammation; and/or rare, small granulomas; and/or glandular epithelial necrosis | Moderate lesions not observed in control animals (affecting 10-20% of the visible surface area): As above with multiple larger foci of mixed inflammation; and/or rare, small granulomas; and/or scattered foci of mineralization; and/or ductular epithelial necrosis |
| **4** (severe) | Widespread moderate to severe mixed inflammation, necrosis, mineralization and/or granulomas (affecting greater than 20% of the visible surface area) | Widespread moderate to severe mixed inflammation, necrosis, ductular rupture, mineralization and/or granulomas (affecting greater than 20% of the visible surface area) | Widespread moderate to severe mixed inflammation, necrosis, mineralization and/or granulomas (affecting greater than 20% of the visible surface area) | Widespread moderate to severe mixed inflammation, necrosis, glandular rupture, mineralization and/or granulomas (affecting greater than 20% of the visible surface area) |
| **5** (severe +) | Same as "4" but with evidence of chronicity such as frequent, extensive mineralization, replacement of necrotic testicular architecture with fibrous connective tissue, or large, well-organized granulomas or abscesses | Same as "4" but with evidence of chronicity such as frequent, extensive mineralization, replacement of necrotic epididymal architecture with fibrous connective tissue, or large, well-organized granulomas or abscesses | Same as "4" but with evidence of chronicity such as replacement of necrotic seminal vesicular architecture with fibrous connective tissue, or large, well-organized granulomas or abscesses | Same as "4" but with evidence of chronicity such as replacement of necrotic prostatic architecture with fibrous connective tissue, or large, well-organized granulomas or abscesses |
